# In vivo co-registered hybrid-contrast imaging by successive photoacoustic tomography and magnetic resonance imaging

**DOI:** 10.1101/2021.03.06.434031

**Authors:** Shuangyang Zhang, Li Qi, Xipan Li, Zhichao Liang, Jian Wu, Shixian Huang, Jiaming Liu, Zhijian Zhuang, Yanqiu Feng, Qianjin Feng, Wufan Chen

## Abstract

Magnetic resonance imaging (MRI) and photoacoustic tomography (PAT) are two advanced imaging modalities that offer two distinct image contrasts: MRI has a multi-parameter contrast mechanism that provides excellent anatomical soft tissue contrast, whereas PAT is capable of mapping tissue physiological metabolism and exogenous contrast agents with optical specificity. Attempts have been made to integrate these two modalities, but rigid and reliable registration of the images for in vivo imaging is still challenging. In this paper, we present a complete hardware-software solution for the successive acquisition and co-registration of PAT and MRI images in *in vivo* animal studies. Based on commercial PAT and MRI scanners, our solution includes a 3D-printed dual-modality animal imaging bed, a 3-D spatial image co-registration algorithm with bi-model markers, and a robust modality switching protocol for in vivo imaging studies. Using the proposed solution, we successfully demonstrated co-registered hybrid-contrast PAT-MRI imaging that simultaneously display multi-scale anatomical, functional and molecular characteristics on healthy and cancerous living mice. Week-long longitudinal dual-modality imaging of tumor development reveals information on size, border, vascular pattern, blood oxygenation, and molecular probe metabolism of the tumor micro-environment at the same time. Additionally, by incorporating soft-tissue information in the co-registered MRI image, we further show that PAT image quality could be enhanced by MRI-guided light fluence correction. The proposed methodology holds the promise for a wide range of pre-clinical research applications that benefit from the PAT-MRI dual-modality image contrast.

## Introduction

Tomographic imaging of living animals has been an important task for preclinical research because it provides cross-sectional images of the subject without surgical intervention. This unique capability has differed it from other transmissive or reflectance imaging approaches such as whole body fluorescence imaging^1^ or digital radiography^2^. Among many tomographic imaging techniques, Photoacoustic Tomography (PAT) and Magnetic Resonance Imaging (MRI) are two advanced biomedical imaging modalities that have been used in various pre-clinical imaging applications ranging from tumor screening^3–7^, therapy evaluation^8–10^, to functional brain imaging^11–16^ and so on. In PAT, an image that maps the original energy deposition inside the target is formed by detecting and processing the ultrasonic signals generated by laser illumination^17, 18^. PAT is able to reveal the distribution of endogenous tissue absorbers, such as oxyhemoglobin (HbO_2_) and deoxyhemoglobin (Hb), and exogenous optical probes, such as the FDA approached Indocyanine Green (ICG), by identifying their absorption spectrum under multiple wavelength excitations^19–21^. On the other hand, as a Nobel winning technology, MRI provides cross-sectional images of the subject by using non-ionizing electromagnetic radiation and measuring the nuclear magnetic resonance signal, and thus enables multitudinous tissue contrast. MRI is able to provide comprehensive, multi-parametric information on anatomy, function and metabolism. Thanks to the emergence of diffusion MRI, functional MRI and other technologies, MRI has covered various clinical neurological, psychiatric, cardiac and abdominal applications^22^.

Given their outstanding imaging capabilities, these two imaging modalities are complementary at multiple dimensions. Firstly, they have distinct image contrast mechanisms. PAT provides molecular imaging capability that reflects the optical characteristics of light absorbers inside tissue^23–25^, whereas MRI images the density of hydrogen protons such that soft-tissue contrast is revealed. Secondly, the imaging speeds are complementary. Thanks to recent advancement in laser technology, PAT imaging speed could reach over 7000 frames per second^26^, whereas high-field MRI system could only acquire at most one to two images per second without scarifying image quality^27^. Thirdly, their spatial resolutions are matched. Common spatial resolutions for commercial pre-clinical MRI and PAT scanners are both around tens to hundreds micrometres^28^ given an imaging field of view of several square centimetres. Finally, PAT and MRI also share the advantages of being non-invasive, non-ionizing and label-free.

The benefits brought about by these complementary features of the two imaging technologies are abundant and attractive. Due to the inherent low soft tissue contrast in PAT, it is difficult to precisely identify the anatomical location of the targeted chromophores. With its combination with MRI, which provides excellent anatomical definition and soft tissue contrast, this limitation might be removed and accurate target lesion positioning and analysis could be achieved. Also, the high imaging speed of PAT could compensate for its counterpart in MRI, making dynamic imaging of transient biological activities such as neuron firing possible. Moreover, information sharing between the acquired dual-modality images might be able to help improve the image quality of one another (e.g. the structural information provided by MRI may be used to guide PAT image reconstruction or image recovery). PAT-MRI dual-modality imaging that simultaneously acquires structural, functional, and molecular images has great potential to push the image analyse focus to multiple scales, allowing for much broader preclinical research impacts.

Previous attempts to integrate images from these two modalities confined to either rigid registration (e.g. imaging of rigid structures such as the brain^29, 30^) or no registration^31–33^. Co-registration of abdomen PAT and MRI images of small animals using a customized single-use silicone MRI holder has been reported previously^34^, which realized the registration of soft tissue images for the first time. Most recently, a prove-of-concept concurrent PAT-MRI imaging system has been proposed with phantom-based feasibility validation^35^. Development of such system requires high cost for customized instrumentation, and sacrifices the flexibility of individual system. Apart from hardware registration, robust software registration algorithms are required to precisely align the images from different modalities. Co-registration of PAT and MRI images of the brain of small animals has been proposed^29^. It combines mutual information based rigid registration algorithm with manually labelled anatomical landmark for the matching of the brain, which is a non-deformed object.

Although there were these aforementioned early demonstrations of successive or concurrent PAT-MRI imaging for various applications, implementing a rigidly co-registered, dual modality imaging solution faced significant challenges. First, PAT imaging requires coupling media (e.g. water) whereas MRI imaging does not. The purpose of the coupling media is to let the excited ultrasound to propagate. However, during modality switching, this coupling media will inevitably affect the posture of the animal. Second, MRI requires a strictly no-metal imaging environment, making the design and fabrication of a robust bimodal registration tool difficult. Third, it requires spatial resolution matching at the axial, radial, and tangential directions, simultaneously. Fourth, flexible, easy-to-use software compensation algorithms for dual-modality image registration are required for high repeatability imaging experiments. Fifth, similar to the attenuation correction in a Positron Emission Tomography/Computed Tomography (PET/CT) scanner where CT image is used to enhance PET imaging, PAT-MRI requires mutual connection and collaboration between the two modalities in order to excavate deeper information. Finally, the obtained dual-model images from long-term in vivo longitudinal imaging should be validated, and the performance of the whole imaging protocol should be benchmarked and analysed.

Here, we propose a method for the successive acquisition and co-registration of PAT and MRI data in *in vivo* mice studies. The method includes a novel dual-modality imaging bed and a robust dual-modality spatial co-registration protocol. The 3D-printed imaging bed is specifically designed to secure the posture and position of the animals during modality switching. Based on this bimodal imaging bed, we design a rigorous data acquisition procedure, a stable modality switching protocol, and a highly automated data post-processing software suite to enable precise matching of the dual-modality images of the entire animal body. We demonstrate the excellent capability of this successive PAT-MRI dual-modality imaging method in *in vivo* applications including tomographic hybrid contrast observation of important organs (simultaneously structural, functional, and molecular imaging), multi-timescale longitudinal monitoring of tumor development (from minute-scale drug uptake to week-long evolution of tumor size and hypoxia condition), and structural MRI guided light fluence correction for quantitative PAT. This integrated and standardized protocol for in vivo small animal PAT-MRI dual-modality imaging will help unlocking and promoting even more preclinical research applications of these two modalities, such as simultaneous functional-anatomical brain imaging, bimodal contrast agent development, and anatomically specific pharmacokinetic research and etc.

## Results

### Spatially co-registered successive PAT-MRI imaging

We first describe the proposed co-registered successive PAT-MRI imaging solution. Our solution is developed particularly for a 7-tesla MRI scanner (Pharmascan, Burker, Germany) and a commercial multispectral cross-sectional PAT system (MSOT inVision128, iThera Medical, Germany), both of which are among the most popular commercial imaging platforms for pre-clinical small animal imaging. The radial resolution of the MRI and PAT systems were similar (~ 150 um). The axial resolution of MRI could reached <100 um, but for the PAT it is limited to 800 um (see METHODS for detail). However, the axial resolution of MRI is tunable such that the spatial resolutions, both axial and radial, are matched for the two modalities. To ensure precise spatial co-registration of the animal during modality switching, our solution includes a specially designed dual-modality imaging bed (Fig. 1 a). This bed consists of a gas tube, a breathing mask, two fixing plates, two ancillary supports and a solid animal support that can be separated into two parts, one for PAT, and the other for MRI. All the components except for the gas tube are 3D printed with polylactic acid (PLA). The gas tube, made of rubber, was connected to the anesthesia gas inlet and the breathing mask to keep the animal under anesthesia. In addition, a silicone sealing pad was used to isolate the frontal part of the mouse head from water during PAT imaging. During the MRI imaging, the MRI support could be fitted into the original MRI animal bed. During the T2 spin echo sequence, the PLA material of the support will not generate interference signal to the object. During PAT imaging, the PAT support was firmly fixed on the original animal holder of the PAT system such that the animal was placed right in the centre of the detector array.

**Fig. 1.**
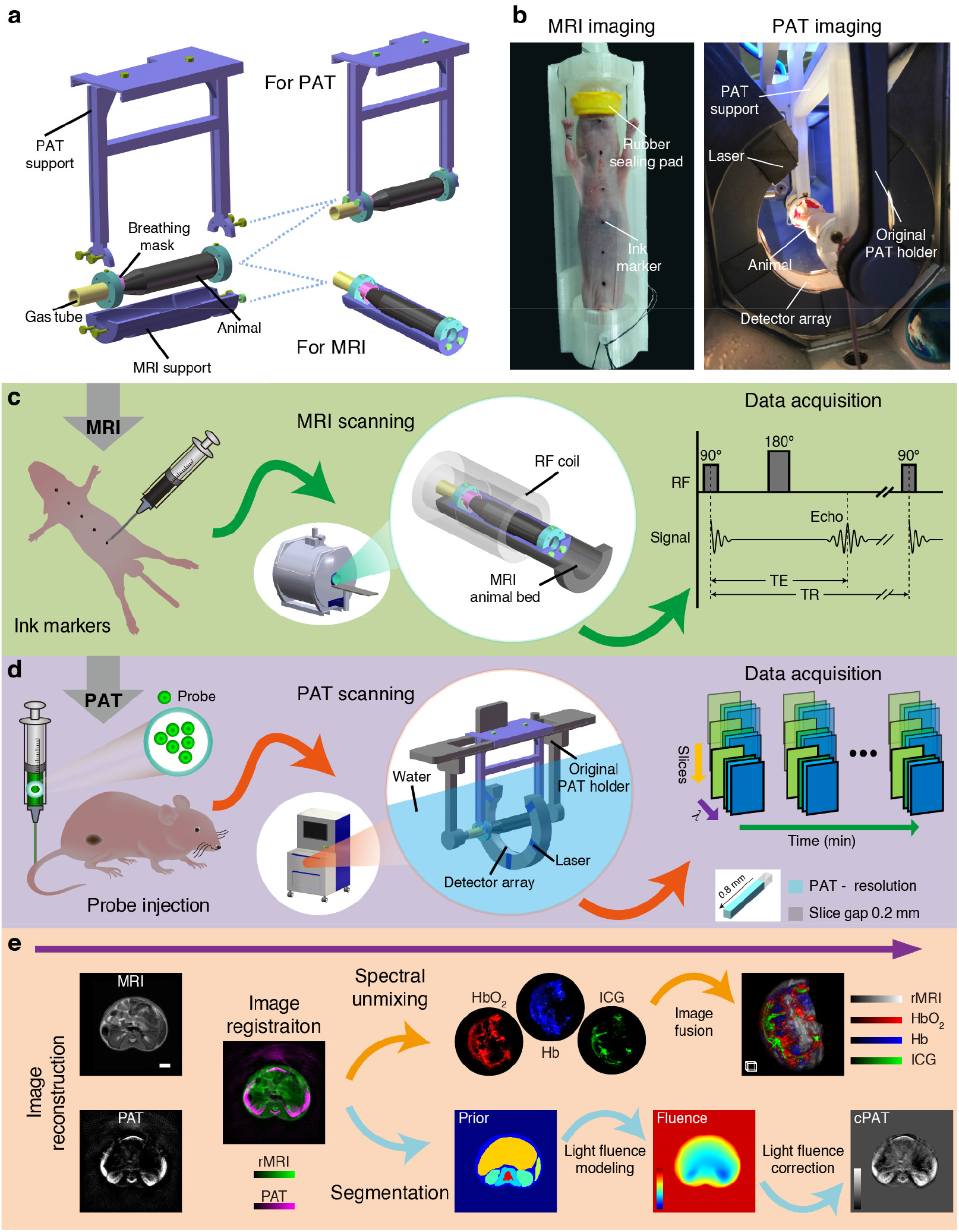
Co-registered hybrid-contrast imaging of living mice by successive PAT and MRI. **a** The dual-modality imaging bed includes a PAT support, a MRI support, a gas tube, a breathing mask, two fixing plates and two ancillary supports. Image acquisition for the two modalities can be realized by assembling different components. **b** Photographs of a nude mouse placed on the animal bed before MRI and during PAT imaging. **c** MRI image acquisition process. Axial marker assignment: the skin surface of nude mice is marked with Chinese ink. MRI scanning: after assembling the MRI support, the animal was placed on the animal bed and transferred to the centre of the magnet for scanning. MRI data acquisition: a 2-D spin echo sequence was used to acquire MRI images. **d** PAT image acquisition process. Probe injection: optional exogenous probe such as ICG was injected through the tail vein. PAT scanning: after assembling the PAT support, the animal was loaded into the PAT imaging chamber filled with water for scanning. PAT data acquisition: 5-D multi-wavelength multi-slice longitudinal PAT images are acquired. **e** Image post-processing process. This process includes image reconstruction, image registration, spectral unmixing, image fusion, and PAT light fluence correction. See text and Supplementary Methods for more detail on image post-processing. Scale bars, 3 mm. Scale box, 1 mm^3^. Details of the dual-modality imaging bed are shown in Supplementary Video 1 and Supplementary figure 1 and 2.

The image acquisition process was divided into four steps (Fig. 1c, d, Supplementary Figure 1). 1) Axial marker assignment: Chinese ink was used as markers for axial registration because it can be visualized in both MRI and PAT. The ink was marked on the skin surface of nude mice using a Gauge 20 needle. The marker size was less than 1 mm and the separation distance was 1 cm such that minimum interference is caused to the images. 2) MRI imaging: the animal was fastened on the MRI support and transferred to the MRI imaging cavity, and the location of the mouse to be scanned was positioned in the center of the RF coil. During MRI imaging, we used a 2-D spin-echo sequence to scan the axial image in the XY plane. After the acquisition was completed, the dataset was resliced along the XZ and YZ directions to obtain the coronal and sagittal images. 3) Support switching: First, we used screws to fasten the MRI support on the fixing plates. Then, we unscrewed the screws on the cantilever and removed the PAT support. During this process, the body of the mouse was always in a tight state that ensured an unchanged posture. After the animal support was switched, exogenous contrast agent for PAT imaging such as ICG could be injected. 4) PAT imaging: we fix the PAT support on the original animal holder of the PAT system, and connect the gas tube to the anaesthesia port on the holder. The whole assembly was then transferred to the imaging chamber filled with distilled water pre-heated to 34-degree Celsius. The assembly was set on a built-in motorized translation stage, such that the animal could be positioned to the optimal field of view. Finally, multi-spectral 3-D PAT image acquisition was carried out. When the PAT imaging section was finished, the PAT support was taken out from the imaging chamber and the animal was released. A video showing the above processes are available in Supplementary Video 1.

After the dual-modality imaging, post-processing of the acquired images were employed to refine the co-registration. The post-processing procedures (Fig. 1e, Supplementary Figure 3) are as follows: 1) PAT image reconstruction: to get rid of the effect of the PAT image background signal on the registration result, we used a non-negative model-based iterative image reconstruction algorithm which limits the signal value to positive during each iteration. 2) Axial registration: the ink markers on the skin surface of nude mice can be visualized on both PAT and MRI, therefore, the corresponding axial position of the images is found by locating the markers (see Supplementary Methods for details). 3) Transverse registration: after the images were axially registered, we performed transverse registration on corresponding PAT-MRI image slices by using a rigid co-registration algorithm based on mutual information. This will compensate for the small shifting or rotation of the animal during modality switching, and further align the dual-modality images. 4) Spectral un-mixing: for PAT images acquired with multispectral excitation, we perform spectral un-mixing to identify the distribution of endogenous (HbO_2_, Hb) or exogenous absorbers (such as ICG) from the background. The un-mixing is based on a linear algorithm^21^ with multispectral PAT images as inputs. 5) Light fluence correction: to account for the light attenuation during its propagation in tissue, we make use of the rich and clear soft-tissue contrast in MRI to guide the estimation of the light fluence distribution during PAT imaging, and design an iterative algorithm (see METHODS for detials) to correct for the light attenuation. The obtained corrected PAT images not only show evenly distributed image intensity, but also represent quantitative optical absorption and scattering coefficients of different tissues.

### Co-registered anatomical imaging by PAT and MRI

With the proposed successive PAT-MRI dual-modality imaging method, we achieved three-dimensional co-registered anatomical imaging of healthy and tumorous mice *in vivo*. Supplementary Fig. 4a shows the corresponding PAT and MRI images at the ink marker position. In the MRI image, the ink marker can be localized easily, however, in the PAT image, the marker spreads for nearly 5 mm along the axial direction (Supplementary Fig. 4b) due to its strong optical absorption, making the identification of the correct PAT slice difficult. To tackle this, we quantified the intensity of the marker at each slice, as plotted in Supplementary Fig. 4c, and then applied Gaussian fitting to the plot. The PAT image at the peak position of the fitted curve was considered to be the correct image that matched the MRI image. Fig. 2a, b show the co-registration results of the healthy and tumorous nude mouse respectively. By resampling and interpolating the XY plane image stack, we obtained the sagittal and coronal images and displayed them in 3D as volume images. The joint visualization of the PAT and MRI images reveals successful matching of the internal structure and the consistency of the body shapes of the animal. The kidneys, spleen and spine in the axial image of healthy nude mice are correctly overlapped, and the body contours are also consistent. The tumorous mice datasets furthered demonstrated the robust performance of the co-registration strategy across week-long continuous dual-modality screening. The bright area around the centre of the tumor in the PAT image, which indicated a decrease of blood oxygen saturation, accurately matched that of the MRI image that showed weaken T2 signal (Supplementary Fig. 5). Finally, we perform quantitative evaluation of the registration performance (Fig. 2c) using the Dice Similarity Coefficient (DSC), which measures the percentage of the overlap between the two images (See Supplementary Methods). The average DSCs for healthy and tumorous mice are 93.06% and 95.12% respectively, demonstrating very high overlap between the PAT and MRI images.

**Fig. 2.**
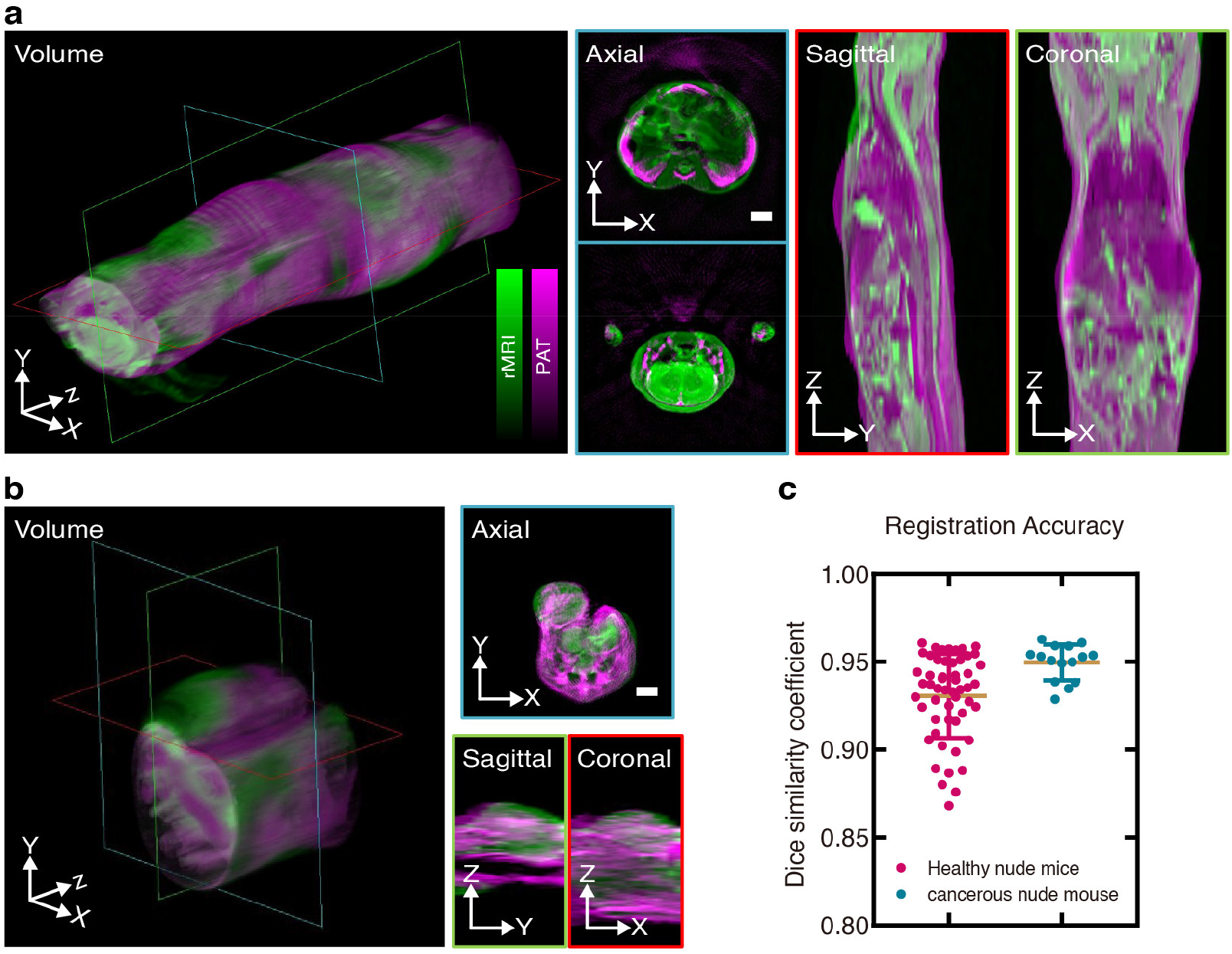
Spatial co-registration results. **a** The registered PAT-MRI dual-modality images from healthy nude mouse. **b** The registered PAT-MRI dual-modality images from tumorous nude mouse. All images in (**a**) and (**b**) are displayed with pseudo-color overlay (magenta for PAT and green for MRI). All sagittal and coronal images are resampled and interpolated from the XY plane image stack, and display in 3D as volume images. **c** Quantification of the co-registration accuracy in Dice similarity coefficient using image data from (**a**) and (**b**). Scale bars, 3 mm.

### Hybrid-contrast longitudinal recording of tumor growth

To demonstrate the capability of simultaneously anatomical and functional imaging, PAT-MRI dual-modality longitudinal observation on nude mice bearing 4T1 tumor was performed. The successive imaging was carried out on day 3, 5, 7, 9, 11, 13 and 17 post tumor implantation. MRI-T2 images that represent structural information and HbO_2_, Hb, HbT images that show hypoxia microenvironment obtained from spectrally un-mixed multispectral PAT images. The image registration results and the distribution of HbO_2_ and Hb separated from the multispectral PAT images are shown in Supplementary Fig. 6. To facilitate better visualization, we used the registered MRI images as a structural priori to manually segment the tumor. As shown in the dual-modality images taken at day 17 (Fig. 3a), the MRI-T2 image showed highly corresponding tumor geometry that matched the distribution of Hb, HbO_2_, and HbT. The white solid-line area depicts the regions with consistent or diametrically opposite features on these images (darker in the T2 and brighter in the Hb and HbT). T2 signal represents changes in blood oxygenation, and weakens when the concentration of Hb inside the tumor increases. To further analysis the correlation between the obtained structural and functional information, profiles (Fig. 3b) of the Hb, HbO_2_, and HbT images were obtained along the white dashed-lines (Fig. 3a) and then compared with that of the MRI image to calculate the Pearson Correlation Coefficient (PCC) (Fig. 3c). The PCC values between T2 and Hb, HbO_2_ and HbT are −0.9069, −0.0048 and −0.9062, respectively. The profiles show opposite spatial trends between T2 and Hb, HbT across the tumor, and the PCC values close to −1 further illustrate the negative correlation of this spatial variation. Overall, the inspection of T2 and Hb revealed the existence of negative correlation, which became higher with time (Supplementary Fig. 6). Moreover, as shown in a series of 3D fusion display using T2, Hb and HbO_2_ images obtained during the tumor development in Fig. 3d, the growth of the tumor was accompanied by the development of neovasculature and the increase of tumor dimension. We also measured the change of tumor volume (Fig. 3e) from the MRI images and calculated the values of tumor oxygen saturation (Fig. 3f) from the distribution of Hb and HbO2. Quantitative parameters obtained from the PAT images indicated that during tumor growth, the Oxygen Saturation (SO_2_) increased continuously from 60.67 % on day 3 to 72.96 % on day 7 for the whole tumor area and decreased from 40.68 % on day 9 to 17.08 % on day 17 for the tumor center alone. The overall volume of the tumor reached a 6-fold increase from 47.73 mm^3^ on day 3 to 339.75 mm^3^ on day 17. In addition, the volume of the central region of the tumor saw a 40-fold increase from 0.44 mm^3^ on day 7 to 17.7 mm^3^ on day 17. This experiment demonstrated the unique capability of label free, long term structural and functional hybrid contrast imaging of the PAT-MRI bimodal imaging method.

**Fig. 3.**
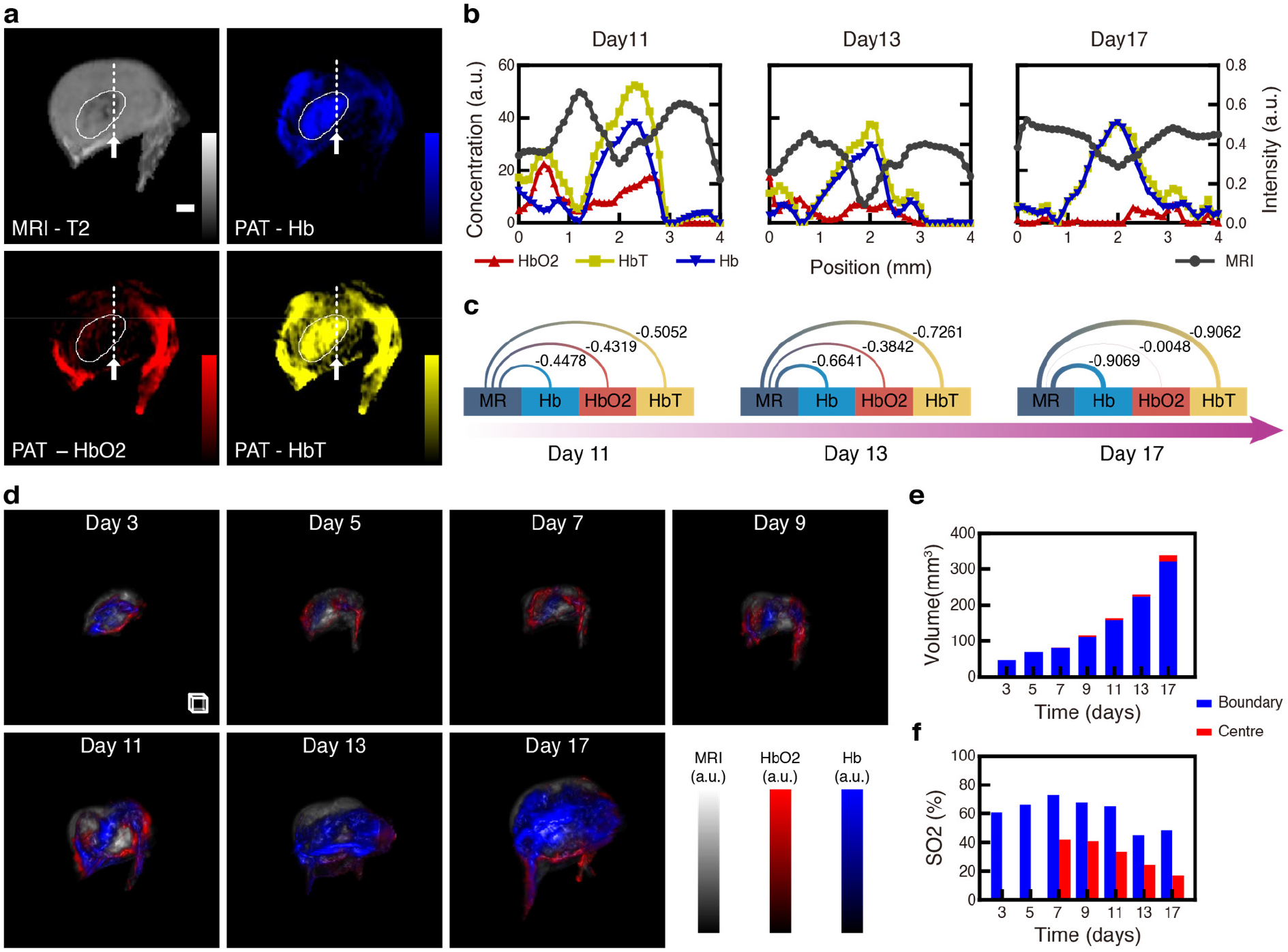
Week-long longitudinal monitoring of tumor dimension and hypoxia microenvironment in living nude mouse by PAT-MRI imaging. **a** MRI image and PAT-derived Hb, HbO_2_ and HbT images in the central XY plane of the 4T1 tumor. The white solid-line area depicts the regions with similar or opposite features. **b** Image profiles drawn along the straight white dashed-lines in (**a**). **c** Pearson correlation coefficients of the profiles between the Hb, HbO_2_, HbT images and the MRI image. **d** 3D fusion display of MRI images and Hb, HbO_2_ images in longitudinal imaging of the 4T1 tumor. We delineated the mask of hypoxic regions inside the tumor and computed the volume size (**e**) from MRI images and the blood oxygen saturation (**f**) from PAT images in the centre and boundary of the tumor separately. Scale bar, 1 mm. Scale box, 1 mm^3^. See also Supplementary Fig. 6 and Supplementary Video 2.

### Spatially localized high-speed imaging of molecular probes

To harness the high speed imaging capability provided by PAT imaging, we applied intravenous (IV) injection of 200 μl of ICG (0.05 μg/μl) on a day-21 4T1 tumorous mouse after MRI imaging, and then performed PAT temporal imaging of the mouse at 5-minute intervals for 40 mins. ICG, which is a FDA-approved NIR fluorochrome commonly used as a contrast agent in retinal and tumor imaging, is able to metabolize in blood-rich organs and excreted into the bile within one hour^36–38^. The registration results and the distribution of ICG identified from the multispectral PAT images are shown in Supplementary Fig. 7. Fig. 4a shows the volumetric images of MRI-T2, HbO_2_, Hb and ICG at various time points. As can be seen, almost immediately after the injection, a small amount of ICG signal appeared in the tumor. Then the signal gradually increased, indicating fast deposition of the ICG inside the tumor. Nevertheless, ICG only appeared in the boundary region of the tumor, and its concentration in the central region was relatively low across the whole imaging period. This indicated a hypoxic area had been developed in the center of the tumor. Furthermore, we measured the ICG concentration at different time at both the centre and boundary regions of the tumor, and the result was shown in Fig. 4b. As can be seen, after ICG injection, there was no significant change in its concentration around tumor centre (< 5 % fluctuation). However, the ICG concentration in the boundary kept on rising around the first 20 minutes and reached a peak increase of 39.12 % at t = 20 minute, and then slowly went down. This preliminary demonstration of spatially localized continuous monitoring of contrast agent reveals the potential of our proposed method for structural enhanced dynamic molecular imaging.

**Fig. 4.**
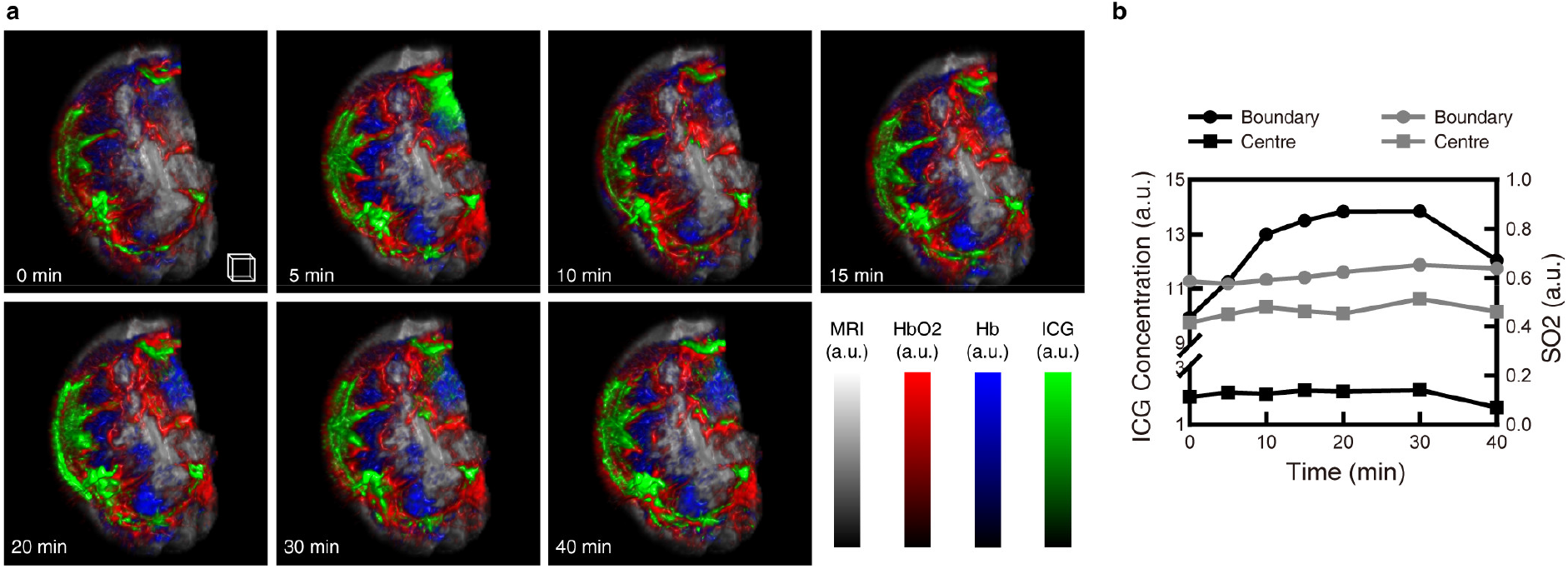
Temporal anatomical-molecular imaging of ICG perfusion inside 4T1 tumor. **a** 3D fusion display of the Hb, HbO_2_, ICG distributions on top of a co-registered MRI image of the 4T1 tumor microenvironment. PAT images were acquired at a 5 minute interval. **b** Time trace of the ICG concentration and SO_2_ of the hypoxic centre and boundary region of the tumor. Scale box, 1 mm^3^. See Supplementary Video 3.

### MRI assisted light fluence correction for quantitative PAT

PAT image is the product of the absorption coefficient and the light fluence distribution^39, 40^. Because light attenuates as it propagates into deeper tissue, to determine the concentration of the chromophores, this light attenuation effect has to be corrected. To eliminate the light fluence from the PAT image and recover the distribution of the optical absorption coefficient is of great significance. Light fluence correction method thatrequired the segmentation of the image at organ level are proposed previously to estimate the optical parameters of each region^40^. However, it is difficult to perform accurate organ segmentation on the PAT images alone because of its poor tissue contrast. Incorporation of co-registered MRI images might be able to solve this problem. Here, we propose MRI structural information guided light flucence estimation and correction for PAT. This method performs segmentation on the registered MRI image acquired with our dual-modality imaging approach, and then uses the segmentation result to guide the estimation of light fluence distribution (see METHODS and Supplementary Fig. 8).

We first performed validation of the method on phantom imaging experiments, and the results are shown in Supplementary Fig. 9. The phantom is a cylindrical tissue mimicking phantom with three rod-shape inclusions, which contained the same material at the same concentration. As can be seen, because of light attenuation, the signal intensity of the inner rod is lower than the outer rods in the uncorrected PAT image. In the corrected image, this attenuation effect has been successfully compensated for using the proposed algorithm. The profiles (Supplementary Fig. 9b) drawn along the three rod shape inclusions show that their photoacoustic signal intensity has returned to the same level after correction.

Next, we applied the proposed light fluence correction algorithm to the *in vivo* healthy nude mice data. Fig. 5a shows the light fluence correction results at the neck and the kidney position, including the raw PAT image (PAT), the registered MRI image (rMRI), the regions segmented on the MRI image (Prior), the light fluence distribution obtained from the optimization algorithm (Fluence), and the corrected PAT image (cPAT). As expected, the light fluence decreases radially from the surface to the center of the animal body. With the application of light fluence correction, the signal around the image centre has been significantly enhanced. And the inner kidney and the outer kidney have achieved similar signal strength compared with the original PAT image. Fig. 5b shows the 3D light fluence distribution obtained by using the proposed MRI-guided light fluence simulation method. As shown in Fig. 5c, after correction, the visibility of the blood vessels deep inside the body have also be enhanced. The proposed dual-modality PAT-MRI imaging strategy made it possible to incorporate the MRI structural information into the light fluence correction process of PAT imaging, and the above experiments have successfully demonstrated the feasibility of this technique.

**Fig. 5.**
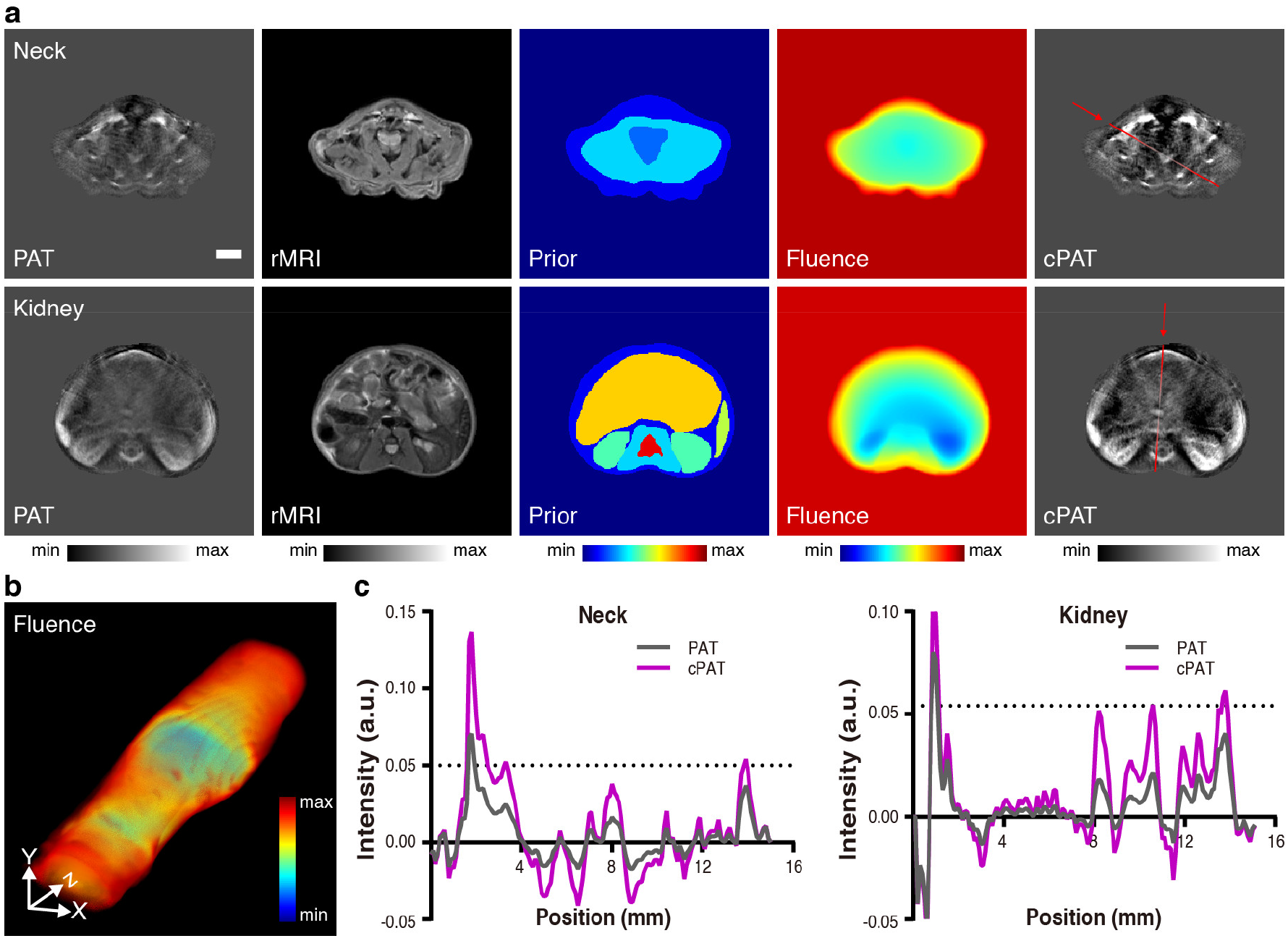
MRI structural guided light fluence correction for nude mouse PAT imaging in vivo. **a** Light fluence correction results at the neck and the kidney position. PAT: raw PAT images. rMRI: co-registered MRI images. Prior: organ-level MRI segmentation results used to guide light fluence estimation. Fluence: light fluence distribution solved by the proposed optimization algorithm based on the Prior image. cPAT: light fluence corrected PAT images. **b** 3D view of light fluence distribution obtained with structural guidance from MRI. **c** Image profiles drawn along the straight red solid lines in (**a**). PAT imaging wavelength: 700 nm. Scale bar, 3 mm. The 3D segmentation results of MRI images and the obtained 3D light fluence distribution are shown in Supplementary Video 4.

## Discussion

Visualization of complementary information derived from different imaging methods provides multiple types of image contrast, such that hidden information of the research problem may be revealed. Imaging performed successively on different modalities saves the flexibility of individual modality, but the imaged animal is easily deformed during modality switching, resulting in anatomy dislocation of the acquired images. This problem makes it difficult to accurately share information provided by the imaging methods because the image contrasts are misaligned. For successive PAT-MRI imaging, on one hand, the structural and functional contrasts, spatial resolutions, and imaging speeds of the two methods are perfectly complementary, making the implementation of such dual-modality imaging attractive. On the other hand, however, since the animal fixation methods for the two modalities are very different, the aforementioned registration problem becomes much more challenging.

The presented work provides a unique strategy for meeting this challenge. To ensure the posture and position of the animal during modality switching, our successive PAT-MRI imaging method involves a hardware registration device and a software toolset with automatic processing capability. On the hardware side, a novel small animal dual-modality imaging bed was designed. This 3D-printed imaging bed solves the water coupling problem in PAT imaging and ensures that the animal does not deform or displace while switching between MRI and PAT. Its introduction preserves the consistency in the shape of the entire mouse body or local lesion contour such as tumor boundary, and thereby simplifies the multi-modality image co-registration problem into a rigid registration problem solvable by standard image processing algorithms. Moreover, animal fastening method of the designed imaging bed was similar to that of the original PAT system. Therefore, PAT imaging artefacts induced by the bed was minimized and animal preparation time was not increased. The use of the breathing mask allowed for the animal to breathe freely underwater during PAT imaging and ensured high survival rate of the animals. Also, assembling and disassembling procedures of the MRI/PAT supports were designed to be both convenience and stable, such that changes in animal pose and position were subtle. The animal bed is simple to manufacture, low cost, reusable, and compatible with the harsh MRI environment. On the software side, an axial registration method based on external dual-modality markers and a transverse co-registration algorithm based on mutual information between MRI and PAT were developed to further improve image co-registration performance. The Chinese ink marked on the animal surface is not only minimum invasive and harmless, but also has PAT-MRI dual contrast. With the above unique advantages, our proposed dual-modality imaging strategy offers a unified, standardized, and convenient solution to implement successive acquisition of PAT and MRI images for *in vivo* preclinical animal research.

Furthermore, the feasibility of the proposed strategy was demonstrated in various dual-modality imaging scenarios. Firstly, healthy nude mouse and cancerous nude mouse spatial co-registration results showed high overlap of animal anatomy on the two images, illustrating the robustness and favourable performance of the proposed strategy. Secondly, dual-modality characterization of spatial and temporal heterogeneities of the hypoxia tumor microenvironment visualized vascular pattern changes throughout the entire tumor development period, revealing the possibility of label-free, multi-contrast monitoring of cancer development. Thirdly, high speed spatial and temporal tomographic imaging of exogenous contrast agent uptake inside tumor demonstrated highly accurate structural localization of the imaging probe, allowing for the study of drug metabolism dynamics with high spatial specificity. Finally, we found that simple manual segmentation on MRI-T2 images provides valuable structural guidance to enhance the estimation of light fluence distribution in PAT imaging. This enabled accurate light fluence correction of the PAT images, and resulted in improvement of the visualization of deeply suited organs and vasculatures.

Based on our work, we envision vast applications by the proposed successive PAT-MRI imaging technology. For example, some studies have shown that the central hypoxia of solid tumors is related to prognosis and treatment resistance^41, 42^. Therefore, longitudinal quantitative analysis of SO2 and HbT with PAT imaging, which is able to access the hypoxic micro-environmental changes, and longitudinal tumor morphology observation with MRI imaging, which can monitoring tumor dimension and growth rate, can work together to provide a platform for the *in vivo* and *ex vivo* evaluation of anticancer therapies aimed at reducing hypoxia and inhibiting tumor angiogenesis^42, 43^.

In this work, we have demonstrated the feasibility of an image acquisition and co-registration method for PAT and MRI. The design of the dual-modality animal imaging bed ensures that the deformation of the animal is within acceptable range when switching imaging modalities, thereby simplifying image co-registration. The dual-modality hybrid-contrast image obtained with our method simultaneously provides functional and structural information. This simple and reliable method can be widely implemented for various PAT-MRI dual-modality *in vivo* animal studies.

## Methods

### PAT-MRI dual-modality imaging bed

The major purpose of the dual-modality animal imaging bed is to ensure that the animal maintains at the same positioning and posture during successive PAT and MRI imaging. Because the animal has to be bathed in water during PAT imaging, the biggest challenge is to make sure the animal position does not changed during the switching between the two imaging modalities. This was achieved by designing the animal bed according to the PAT and MRI system environment and geometric dimensions. The dual-modality imaging bed can be separated into two parts, one for PAT, and the other for MRI. All the components except the gas tube were 3D-printed with PLA material using a desktop 3D printer (Jenny3D, China).

#### Gas tube

A 10 mm diameter tube that connects the anaesthetic port to the breathing mask. The gas tube is made of rubber and therefore it can supply the anaesthetic gas to the animal while preventing water from entering the mask.

#### Breathing mask

The breathing mask functions like a swimming goggle except that it only covers the mouth and nose of the animal. It includes a funnel-shaped structure, a built-in copper wire, and a sealing silicone pad. One small end of the mask is connected to the anaesthetic port through the gas tube. The other end of the mask is sealed with a silicone pad with a small hole in the center, such that the frontal part of the mouth head fits into the hole. The copper wire is mounted transversely inside the breathing mask for hooking the teeth of the mouse during PAT imaging. In this way, the mouse face can be closely fitted to the silicone sealing pad, and drowning of the animal can be prevented. The mask also helps to keep the mouse head steady during imaging.

#### Fixing plates and ancillary supports

Both the left and right sides of the imaging bed contained the fixing plate and the ancillary support. The functions of the two fixing plates include: 1) to fix the animal onto the PAT and MRI supports, and 2) to bind the limbs of the mouse and fix the breathing mask. These fixing plates can be firmly attached to the imaging supports by using plastic or copper screws, and then the arms and legs of the animal can also be tied to the support. The two ancillary supports are used to support the torso of the mouse. The interested imaging regions can be selected by simply adjusting the positions of the two ancillary supports. Supplementary Fig. 2 demonstrates four types of fixing plates and ancillary supports for the imaging of different parts of the animal body.

#### PAT support

It includes two components: the mounting plate and the cantilever, which were assembled by screw combination. The mounting plate can be perfectly attached on the original animal holder of the PAT system, and can be translated in the axial direction of the nude mouse (less than 1cm) to facilitate the connection of the gas tube. The cantilever was fastened to the fixing plates by screws such that the animal to be imaged is in a suspended state. This design centers the animal in the ring-shape detector, and lets none obstruction appeared along the sound propagation path. In this way, the PAT image quality can be ensured.

#### MRI support

A curved base-plate used to fix the animal during MRI imaging by screwing to the fixing plates. The MRI support is adapted to the body contour of the animal, prevents the animal from deformation, and matches the shape of the original MRI bed.

### Spatial co-registration algorithm of the PAT and MRI images

The spatial co-registration algorithm of the PAT and MRI images contains the following steps. Step 1) image pre-processing: perform image reconstruction, denoising, and background removal on the collected multi-modal data. Step 2) axial registration with external markers: locate the corresponding position of the multi-modality cross-sectional images by Gaussian fitting. Step 3) 2D transverse co-registration: register the 2D bi-modal images with an automated rigid transverse registration algorithm. Evaluation of co-registration accuracy by calculating the DSC was performed after the above software co-registration was done. The schematic of the proposed spatial co-registration algorithm is shown in Supplementary Fig. 3.

### Light fluence estimation and correction method

The PAT images are formed by reconstructing the original point source of the ultrasonic waves generated by absorbing the laser pulse, and the pixel value 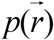 in the image is expressed as:

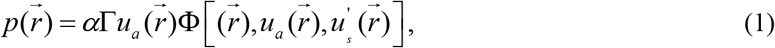

where *α* denotes the system response, Γ denotes the thermo-elastic Gruneisen parameter, 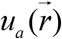 and 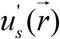 denotes the absorption coefficient and the scattering coefficient, Φ denotes the light fluence within a voxel at the position 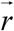. To perform light fluence correction, we first simulate the light fluence distribution Φ over the imaging field-of-view, and then calculate the fluence corrected PAT image by Φ using the following equation:

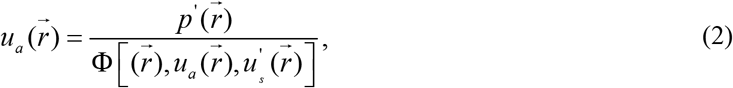

where we make the reasonable assumption that 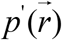 has been reconstructed from the acoustic measurements accurately and with negligible structural distortion^17, 18^.

To make use of the structural information provided by the MRI images, we designed and implemented an iterative optimization method to calculate the absorption and scattering coefficients. The schematic of the algorithm is as shown in Supplementary Fig. 8. To improve the convergence speed of the absorption coefficient *u_a_* optimization, we manually segmented the registered MRI images into different organ regions *R_N_*. These regions exhibited small changes in hemoglobin content and oxygen saturation, and therefore the optical parameters could be considered uniform within an individual region. The segmentation result was later used as prior information for light fluence estimation. We assigned specifically one absorption coefficient and one scattering coefficient to each region according to reference^40, 44^ and then use this as a constraint to solve the light fluence simulation problem. In the optimization algorithm, the light fluence distribution was modelled by the diffusion equation, and solved by using the Toast++ toolbox^45^ in MATLAB (Mathworks, US). The finite element method (FEM) was employed in the Toast++ toolbox to model the light transport inside the object and the optimization of the objective function was implemented with the built-in minimization function ‘fmincon’ in MATLAB. For initialization, the absorption coefficient varied from 0 mm^−^ to 1 mm^−^, and the scattering coefficient was limited to be within a ± 10% variation range.

### Animal models

All animal experiments were approved by the local Animal Ethics Committee of Southern Medical University and were performed in accordance with current guidelines. In the *in vivo* animal imaging experiment, 6 healthy nude mice (12-15 g/each, female, Southern Medical University, Guangzhou, China), and 4 nude mice carrying 4T1 mammary carcinoma (Southern Medical University Cancer Institute, Guangzhou, China) were used. Animals were kept in ventilated cages inside a temperature-controlled room, under a 12-h dark/light cycle. In order to reduce abdominal peristalsis artifacts caused by food digestion and to prevent the mice from excreting and polluting the imaging environment during PAT imaging, the nude mice were fasted for 8 hours before imaging.

### Magnetic resonance imaging

All MRI scan were performed on a 7 T small animal MRI system (Pharmascan, Bruker, Germany) operated by ParaVision 6.0 software platform. A 1H transmit-receive volume coil with 40 mm inner diameter was used for signal transmitting and receiving. The animal was anesthetized with 4 % chloral hydrate at 0.01 ml/g. We assembled the MRI support, fixed the limbs of the anesthetized animal on the fixing plates on both sides, and hooked the teeth of the nude mouse on the copper wire in the breathing mask. In order to reduce the image artefacts caused by respiratory movements, medical oxygen mixed with high concentration isoflurane (1%, RWD, China) was transmitted through the gas tube to the breathing mask, so that the respiratory rate of nude mice was maintained at 15-20 times/min. The body temperature of the nude mouse was monitored using the rectal probe of a small animal monitoring system (Model 1030, Small Animal Instruments Inc., New York, NY, USA), and stabilized at 37 ± 0.1 °C using the heater module. The T2 MRI images of nude mouse were obtained using a 2-D spin echo sequence (Turbo rapid acquisition with refocused echoes) with the following imaging parameters: RARE factor 8, echo time 10 ms, repetition time 6000 ms, 5 averages, slice thickness/gap 0.8/0.2 mm, field of view 25 × 25 mm^2^, matrix 250 × 250, spatial resolution 0.1 × 0.1 × 0.8 mm^3^. Sagittal T2 images of the YZ plane where the markers located were firstly acquired. Based on these images, the slice direction and position of the axial image was selected so that each marker was at the centre of the slice and the slice direction was perpendicular to the long-axis of the animal. The axial T2 images covering either the upper or lower parts of nude mouse were then acquired because the coil’s effective imaging range was insufficient to cover the entire mouse.

### Photoacoustic tomography imaging

A commercial small animal multispectral photoacoustic tomography system (MSOT inVision128, iThera Medical, Germany, Fig. 1d) was employed to perform all the PAT imaging. A pulsed OPO laser (670 nm −960 nm tunable) with pulse width <10 ns, repetition rate of 10 Hz and a peak pulse energy of 60 mJ at 760 nm is employed in this PAT system. The laser light excites the sample through a ten-arm fiber bundle, which provides a diffused, homogeneous, and 360-degree illumination over the surface of the animal. The generated ultrasonic waves are detected by 128 toroidally focused ultrasound transducers with a centre frequency of 5 MHz (60% bandwidth) arranged over an azimuth span of 270-degrees around the cylinder with a radius of curvature of 41 mm (Fig. 1d).

When switching to PAT imaging, we needed to assemble the PAT support on the fixing plates with screws first, and then removed the MRI support. This is to ensure that the posture of the animal did not changed during the modality switching process. Finally, the animal-fixed PAT support was attached to the original PAT holder which aligned it with the center of the transducer and immersed the animal in 34 ℃ heated water for ultrasonic coupling and warm keeping. All nude mice were anesthetized with 1% isoflurane during imaging. Whole body imaging was realized by translating the PAT support axially with the built-in motorized translation stage. For contrast enhanced PAT, an insulin injection needle was embedded into the tail vein in advance, and was connected to a long Polyethylene Tubing 10 (PE 10) that enabled contrast agents injection (such as ICG) from outside the imaging chamber. Multispectral PAT images were acquired at five different illumination wavelengths: 700, 730, 760, 800 and 850 nm and averaged with signals from 10 frames per wavelength. The ultrasound time series signals are then reconstructed into 2D pressure maps using a model-based iterative reconstruction algorithm with a 30 × 30 mm field-of-view at 300× 300 pixels. During image reconstruction, the pressure value is limited to positive because the negative value only reflects the artifacts caused by the incomplete geometry of the system. Finally, the linear unmixing algorithm^21^ was used to calculate the distribution of HbO_2_, Hb and ICG.

### Statistical analysis

Matlab (MathWorks Inc.) and GraphPad Prism software (GraphPad Software Inc.) were used for statistical analyses and graph drawing.

## Data Availability

The data that support the findings of this study are available from the corresponding authors upon reasonable request.

## Acknowledgements

This work was supported by National Natural Science Foundation of China (31700857), Guangzhou Science and Technology Program (201804010375), Pearl River Talented Young Scholar Program (2017GC010282) and Guangdong Provincial Key Area R&D Program (2018B030333001).

